# The mIAA7 degron improves auxin-mediated degradation in *C. elegans*

**DOI:** 10.1101/2022.05.31.494192

**Authors:** Jorian J. Sepers, Noud H. M. Verstappen, An A. Vo, James Matthew Ragle, Suzan Ruijtenberg, Jordan D. Ward, Mike Boxem

## Abstract

Auxin-inducible degradation (AID) is a powerful tool for the targeted degradation of proteins with spatiotemporal control. One limitation of the AID system is that not all proteins are degraded efficiently. Here, we demonstrate that an alternative degron sequence, termed mIAA7, improves the efficiency of degradation in *C. elegans*, as previously reported in human cells. We tested depletion of a series of proteins with various sub-cellular localizations in different tissue types and found that the use of the mIAA7 degron resulted in faster depletion kinetics for five out of six proteins tested. The exception was the nuclear protein HIS-72, which was depleted with similar efficiency as with the conventional AID* degron sequence. The mIAA7 degron also increased the leaky degradation for two of the tested proteins. To overcome this problem, we combined the mIAA7 degron with the *C. elegans* AID2 system (*C*.*e*.AID2), which resulted in complete protein depletion without detectable leaky degradation. Finally, we show that degradation of ERM-1, a highly stable protein that is challenging to deplete, could be improved further by using multiple mIAA7 degrons. Taken together, the mIAA7 degron further increases the power and applicability of the AID system. To facilitate the generation of mIAA7-tagged proteins using CRISPR/Cas9 genome engineering, we generated a toolkit of plasmids for the generation of dsDNA repair templates by PCR.

## Introduction

Perturbing the functioning of proteins is essential to decipher their biological roles, and can be accomplished using a wide variety of approaches (Housden *et al*., 2017). The desire to investigate the functioning of proteins in specific cell-or tissue types or at specific developmental stages has led to the development of several methods that offer spatial or temporal control over protein expression. In *C. elegans*, tissue-specific RNA interference (RNAi) and conditional loss-of-function alleles utilizing targeted nucleases or recombinase strategies can conditionally interfere with protein functioning at the level of the corresponding mRNA or genetic locus (Cheng *et al*., 2013; Davis *et al*., 2008; Hoier *et al*., 2000; Muñoz-Jiménez *et al*., 2017; Qadota *et al*., 2007; Ruijtenberg and van den Heuvel, 2015; Shen *et al*., 2014). To target proteins directly, several conditional protein depletion systems have been developed that mark proteins of interest for degradation by the 26S proteasome through ubiquitination (Armenti *et al*., 2014; Cho *et al*., 2013; Wang *et al*., 2017; Wu *et al*., 2017). The best-known of these systems are the ZF1-mediated degradation, nanobody-mediated degradation, and auxin-inducible degradation (AID) systems. ZF1-mediated degradation and nanobody-mediated degradation both repurpose the endogenous E3 ubiquitin ligase substrate-recognition subunit ZIF-1 (Armenti *et al*., 2014; Wang *et al*., 2017). The ZF1-mediated degradation system targets proteins that are tagged with the ZF1 zinc-finger domain by ectopic expression of ZIF-1, and can be combined with a *zif-1* loss-of-function background to avoid undesired degradation from endogenously expressed ZIF-1 (Armenti *et al*., 2014; Sallee *et al*., 2021). Nanobody-mediated degradation targets proteins that are tagged with a fluorescent protein by exogenously expressing ZIF-1 fused to a nanobody that targets the fluorescent protein (Wang *et al*., 2017). Finally, in the AID system, a protein of interest is tagged with an AID degron sequence derived from Indole-3-acetic acid (IAA) proteins, and combined with the expression of the plant-derived F-box protein transport inhibitor response 1 (TIR1) (Nishimura *et al*., 2009; Zhang *et al*., 2015). TIR1 forms a functional SKP-1– Cullin–F-box protein (SCF) E3 ubiquitin ligase complex with endogenous SCF proteins. In the presence of auxin (indole-3-acetic acid, or IAA), TIR1 associates with the AID degron resulting in poly-ubiquitination of the target protein (Nishimura *et al*., 2009; Zhang *et al*., 2015).

Most of the systems described above allow for spatial or temporal control, but not both simultaneously, by placing components of the system under control of tissue specific or inducible promotors. Combined spatial and temporal control could, however, be accomplished by incorporating bipartite expression systems (Nance and Frøkjör-Jensen, 2019). In contrast, in the AID system, expression of TIR1 from tissue-specific promoters confers spatial control while timing the addition of auxin offers temporal control over protein degradation.

The AID system has rapidly gained in popularity within the *C. elegans* community and various resources to facilitate its usage have been generated. For example, several strains are available that express TIR1 from different tissue-specific promoters (e.g. Ashley *et al*., 2021; Zhang *et al*., 2015) and cloning vectors are available that facilitate the generation of repair templates for genome editing (Ashley *et al*., 2021; Kroll *et al*., 2021; Negishi *et al*., 2022). Several improvements or alterations to the AID system have also been generated. These include synthetic auxin analogs that may cause less cytotoxicity or are more soluble in aqueous buffers (Martinez *et al*., 2020), and an auxin-modification that is more effective at promoting protein degradation in embryos, presumably by better permeating the eggshell (Negishi *et al*., 2019).

Most recently, an improved AID system (AID2) was adapted for *C. elegans* that addresses two caveats of AID: leaky degradation that is observed for a subset of proteins, and impact on the physiology of *C. elegans* caused by exposure to auxin (Hills-Muckey *et al*., 2022; Negishi *et al*., 2022; Yesbolatova *et al*., 2020). Ideally, auxin-induced protein degradation should occur strictly upon the addition of auxin, and the addition of auxin should not cause effects other than the degradation of the target protein. However, the AID system does suffer from leaky or basal degradation of at least a subset of target proteins in the absence of auxin, which can result in undesired phenotypes (Hills-Muckey *et al*., 2022; Li *et al*., 2019; Martinez *et al*., 2020; Mendoza-Ochoa *et al*., 2019; Natsume *et al*., 2016; Negishi *et al*., 2022; Sathyan *et al*., 2019; Schiksnis *et al*., 2020; Yesbolatova *et al*., 2019; Yesbolatova *et al*., 2020). In addition, the millimolar concentrations of auxin needed for maximum degradation efficacy increase the lifespan of worms and activate unfolded protein response pathways resulting in an enhanced protection against ER stress (Bhoi *et al*., 2021; Hills-Muckey *et al*., 2022; Loose and Ghazi, 2021). The improved AID2 system relies on a TIR1 mutation (F79G in Arabidopsis TIR1) that alters the auxin-binding interface to fit the bulky auxin derivative 5-phenyl-indole-3-acetic acid (5-Ph-IAA) (Hills-Muckey *et al*., 2022; Negishi *et al*., 2022). The use of this TIR1 variant greatly reduces leaky degradation, while requiring low micromolar concentrations of 5-Ph-IAA that are less likely to affect the physiology of exposed animals.

One caveat of the AID system that has not yet been addressed in *C. elegans* is that some target proteins demonstrate slow degradation kinetics, incomplete depletion, or both, complicating their functional analysis (Duong *et al*., 2020; Patel and Hobert, 2017; Riga *et al*., 2021; Serrano-Saiz *et al*., 2018). Here, we adapt an alternative AID degron sequence (mIAA7) that was shown to result in faster and more complete protein degradation in cultured human cells for use in *C. elegans* (Li *et al*., 2019). We show that the mIAA7 degron improves degradation for a panel of proteins with different subcellular locations, expressed across different tissues. While the mIAA7 degron did enhance the leaky degradation of two out of four proteins tested, we show that it can be effectively combined with the *C. elegans* AID2 system (*C*.*e*.AID2) to achieve more complete protein depletion without detectable leaky degradation. Finally, we demonstrate that using two or more degrons could improve degradation of ERM-1, the most challenging protein to degrade that we have encountered to date. Thus, the mIAA7 degron further expands the usefulness of the AID system to determine the biological functioning of proteins in *C. elegans*.

## Results

### The mIAA7 degron improves AID mediated degradation

AID degrons are derived from IAA proteins, and consist of the conserved domain required for TIR1 recognition (Domain II) and flanking sequences (Gray *et al*., 2001; Ramos *et al*., 2001). The two most commonly used degron sequences, termed AID* and mAID, are based on the IAA17 protein (Fig. 1A) (Kubota *et al*., 2013; Morawska and Ulrich, 2013). An alternative degron sequence derived from IAA7 was recently shown to result in faster and more complete protein degradation in cultured human cells, compared to the mAID degron (Li *et al*., 2019). This degron, termed mIAA7, contains the IAA7 Domain II and includes a longer N-terminal flanking region than AID* or mAID (Fig. 1A).

**Figure 1.**
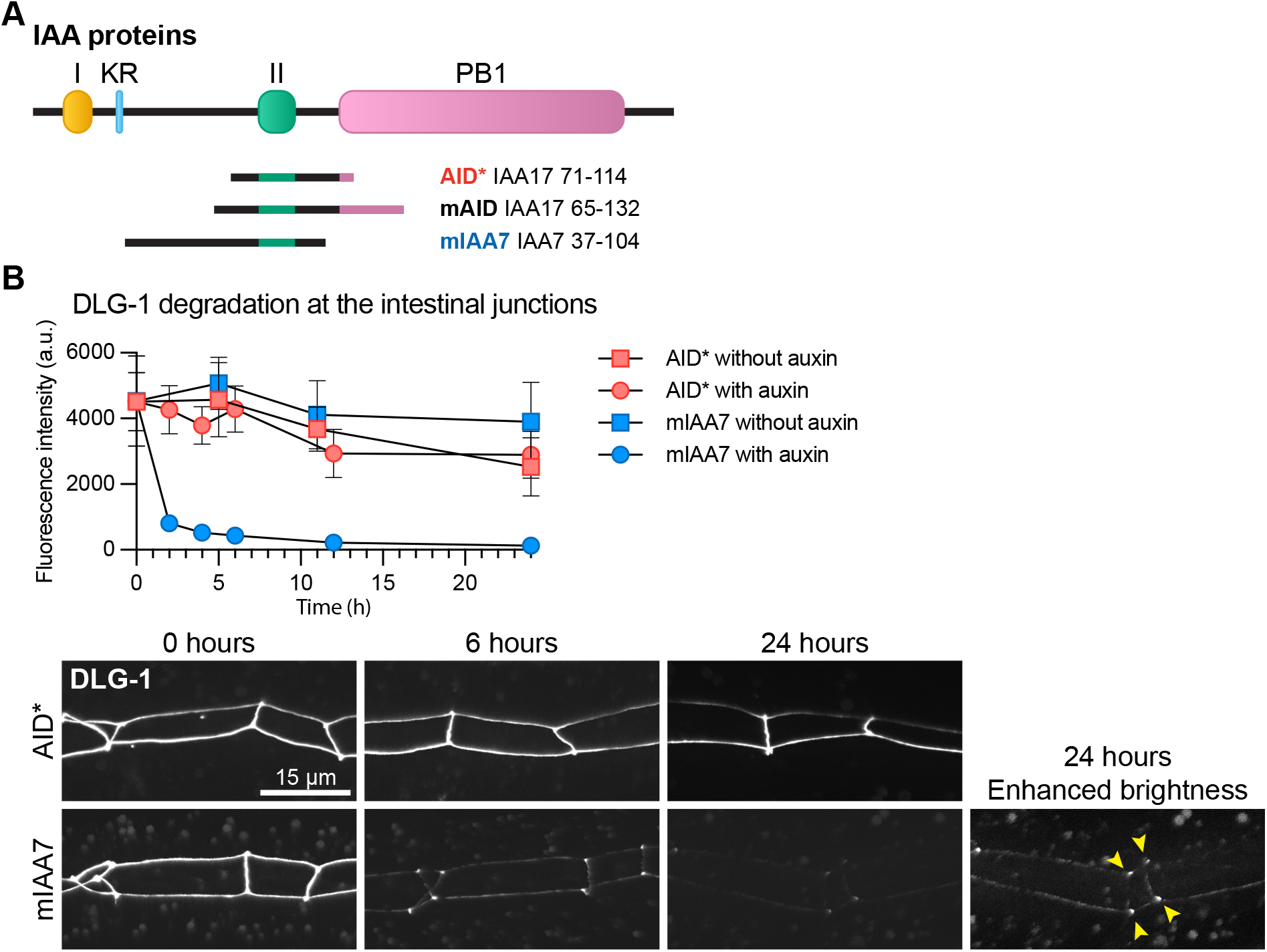
The mIAA7 degron improves auxin-mediated degradation of DLG-1. **(A)** Schematic overview of IAA proteins and the different AID degrons that have been derived from them. IAA = Indole-3-acetic acid; AID = auxin inducible degron; I = domain I; KR = conserved lysine and arginine residue; II = domain II; PB1 = Phox and Bem1p domain. **(B)** Comparison between AID*-and mIAA7-mediated degradation kinetics of intestinal DLG-1 in L3 larvae using 1 mM auxin. Values in the graph are arbitrary units (a.u.) and each data point represents the average intensity at intestinal cell junctions for the given condition and timepoint. Error bars: mean ± SD; n: 7–20 animals. Images shown are representative maximum intensity projections that were acquired and displayed with the same settings for comparison, except for the indicated panel with enhanced brightness to show residual DLG-1. Yellow arrowheads point to the apical junctional sites where three intestinal cells meet.

To assess the effectiveness of the mIAA7 degron in *C. elegans*, we selected the Discs large ortholog DLG-1 as an initial test protein to degrade. DLG-1 is a cell polarity regulator that is involved in the formation of cell-cell junctions and is essential for embryonic development (Bossinger *et al*., 2001; Firestein and Rongo, 2001; Köppen *et al*., 2001; McMahon *et al*., 2001). Previously, we found that the conventional AID* degron was ineffective at mediating degradation of DLG-1 (Riga *et al*., 2021). To determine if use of the mIAA7 degron results in more effective degradation, we tagged the endogenous *dlg-1* locus with codon optimized mIAA7 and GFP sequences by homology-directed repair of CRISPR/Cas9-induced DNA double-strand breaks, using the same insertion site and tag order used previously to generate the *dlg-1::AID*::GFP* locus. We then compared the degradation kinetics of AID* and mIAA7 tagged DLG-1 in the intestine, using a single-copy integrated transgene expressing TIR1 from the intestine specific *elt-2* promoter. We exposed synchronized larvae to auxin from the L3 stage and quantified the levels of DLG-1 at apical junctions of intestinal cells at various timepoints over a 24-hour time-period. Consistent with our previous observations, depletion of DLG-1 tagged with AID* was inefficient, with junctional fluorescence levels similar to controls not exposed to auxin for the complete duration of the experiment (Fig. 1B). In contrast, DLG-1 tagged with mIAA7 was depleted to 4.8% of starting levels after 12 hours of auxin exposure (Fig. 1B). Degradation was not fully complete, as low levels of DLG-1 remained detectable even after 24 hours of exposure to auxin, particularly at the apical junctional sites where three intestinal cells meet (Fig. 1B). Nevertheless, the use of the mIAA7 degron dramatically improved the efficiency of auxin-mediated degradation of DLG-1 in *C. elegans*.

To evaluate the effectiveness of the mIAA7 degron more systematically, we next investigated the degradation kinetics of a variety of proteins using both AID* and mIAA7. The proteins were chosen to represent distinct subcellular localizations and their degradation was investigated in different tissue types. We investigated degradation of the apical polarity regulator PAR-6 in larval seam cells, of the intermediate filament regulator BBLN-1 in intestinal cells, of the adhesion protein SAX-7 in the ALM neurons, and of the ribosomal protein RPS-26 in body wall muscle cells. For each of these proteins, endogenous lines expressing the protein tagged with AID* and a fluorescent protein had previously been generated in our groups. We generated mIAA7-tagged variants keeping the tag order and orientation (N-or C-terminal) and introducing as few other changes to the protein amino acid sequence as possible (Supplementary File S1). We then performed timecourse experiments with synchronized animals to compare the degradation efficiencies, adding auxin at the L2-(PAR-6 and RPS-26) or L3 stage (BBLN-1 and SAX-7) and quantifying fluorescence levels at indicated intervals (Fig. 2A). PAR-6 levels were measured at seam– hyp7 junctions, BBLN-1 levels at the apical intermediate filament layer, SAX-7 levels at the cell wall of the ALM cell body, and RPS-26 in the cytoplasm of body wall muscle cells. PAR-6, BBLN-1, and SAX-7 are already highly efficiently degraded using the AID* degron. For these proteins, we lowered the auxin concentration from 1mM to 5 or 50 μM to increase the time needed for degradation. Under these conditions, for each of these proteins, maximum depletion was achieved twice as fast with the mIAA7 degron compared to the AID* degron (Fig. 2A, S1A). More strikingly, mIAA7-mediated degradation of the ribosomal protein RPS-26 in the body wall muscle cells resulted in complete degradation in 6 hours compared to 24 hours with the AID* degron (Fig. 2A, S1A). Thus, for all four proteins, use of mIAA7 increases the degradation efficiency.

**Figure 2.**
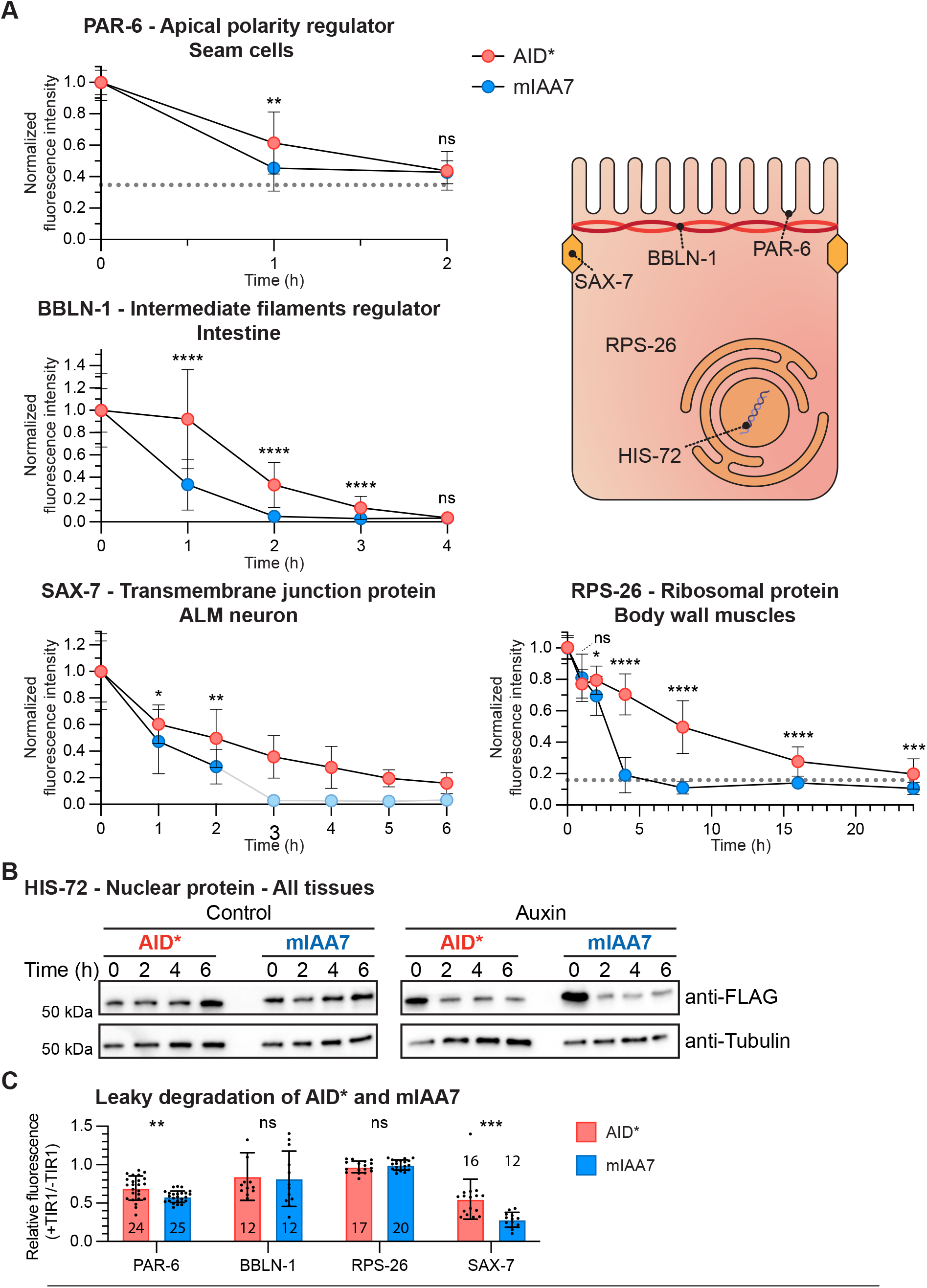
The mIAA7 degron robustly increases the efficiency of AID-induced protein degradation for several proteins across multiple tissues and cellular compartments. **(A)** Comparison between AID*-and mIAA7-mediated degradation for indicated proteins and tissues. Drawing represents a generic polarized cell with the subcellular localization of the proteins investigated indicated. PAR-6 was measured at the apical domain of seam cells in L2 larvae on 5 μM auxin, BBLN-1 was measured at the apical domain in the intestine of L3 larvae treated with 50 μM auxin, SAX-7 was measured at the plasma membrane in the ALM neuron cell body of L3 larvae treated with 5 μM auxin, and RPS-26 was measured in the cytoplasm of body wall muscles of L2 larvae treated with 1 mM auxin. Values are normalized to the mean intensity levels at 0 hours of auxin exposure, and each data point represents the average intensity for the given condition and timepoint. Due to the nature of the quantification for PAR-6 and RPS-26, the fluorescence levels did not reach zero as they did for the other targets. Non-zero baseline level values of wildtype animals are indicated, resembling a completely degraded protein pool. For SAX-7 degradation using the mIAA7 degron, exposure to auxin of 3 hrs or longer resulted in complete depletion and an inability to locate the ALM cell body. For these timepoints, values were plotted at 0 and no statistics were performed (light blue dots). Error bars: mean ± SD; Statistical test: Mann-Whitney U test; n = 9-29 animals. **(B)** Western blots detecting HIS-72:AID*::3xFLAG and HIS-72::mIAA7::3xFLAG in synchronized control or 4 mM auxin-treated L1 larvae. An anti-alpha-tubulin loading control and protein size standard markers in kilodaltons (kDa) are provided. Time in hours (hr) in control media or 4mM auxin containing media is indicated. **(C)** Comparison of leaky degradation between AID* and mIAA7 degrons for the indicated proteins. Measurements for each protein were done as in panel A. Values are normalized to the mean intensity levels of animals expressing the same degron tagged protein but not expressing TIR1. Each data point in the graph represents a single animal. Error bars: mean ± SD; Statistical test: Mann-Whitney U test; n values are indicated in or above the bars.

Finally, we tested the ability of the IAA7 degron to improve the degradation of the nuclear histone protein HIS-72, as several nuclear proteins showed only a modest increase in degradation efficiency with the mIAA7 degron in human cells (Li *et al*., 2019). We previously observed that the degradation efficiency of HIS-72 tagged with AID* varied between cell types, making this protein a good test case to detect a potential difference in degradation kinetics (our unpublished data). To be able to monitor HIS-72 levels in individual cells by immunofluorescence as well as measure overall HIS-72 levels by Western blot, we inserted sequences encoding GFP, the AID* or mIAA7 degron, and a triple FLAG tag at the start codon of the *his-72* locus. We exposed synchronized L1 animals ubiquitously expressing TIR1 to auxin and analyzed HIS-72 levels by fluorescence microscopy and Western blot analysis over a 6-hour period at 2-hour intervals. Control animals were not exposed to auxin. By Western blot analysis, HIS-72 levels were reduced but not absent after 2 hours of exposure to auxin using both the AID* and mIAA7 degrons. HIS-72 levels did not decrease further upon prolonged auxin exposure and using either degron. These findings were corroborated by the fluorescence microscopy analysis. At 2 hours of auxin exposure, HIS-72 levels in cells in the central region of the body were sharply reduced, while HIS-72 levels in cells in the head and tail regions appeared unaffected (Fig. S2B). Further exposure to auxin up to 6 hours did not appear to decrease HIS-72 levels in the head and tail regions. Thus, the mIAA7 degron does not appear to improve the efficiency of HIS-72 depletion, but does function comparably to the AID* degron for this protein.

Taken together, our data demonstrate that the mIAA7 degron robustly increases the efficiency of AID-induced protein degradation across multiple tissues and cellular compartments, except for the nuclear protein HIS-72. The latter result might suggest that the degradation speed of nuclear proteins is mainly limited by the accessibility of TIR1 to the nuclear proteins rather than the degron sequence, though more nuclear proteins will need to be tested to assess the generalizability of this hypothesis. However, this assertion is in line with mIAA7-mediated degradation data in human cells, where improved degradation of nuclear proteins was achieved by expressing TIR1 with a nuclear localization signal (NLS) sequence (Li *et al*., 2019).

### mIAA7 can increase leaky degradation of target proteins

The increase in degradation efficiency conferred by the mIAA7 degron raises the possibility that leaky degradation levels are similarly increased. To investigate this possibility, we determined depletion levels of PAR-6, BBLN-1, SAX-7, and RPS-26 tagged with the AID* or mIAA7 degron as above in strains lacking or expressing TIR1. For RPS-26, we did not observe significant leaky degradation with either degron (Fig. 2B, S1B). For BBLN-1, we observed moderate leaky degradation, with a ∼17% decrease in fluorescence intensity using the AID* degron upon TIR1 expression in the intestine (Fig. 2B, S1B). However, use of the mIAA7 degron did not increase the leaky degradation of BBLN-1 (Fig. 2B, S1B). For PAR-6 and SAX-7, we observed more severe levels of leaky degradation that were significantly increased using the mIAA7 degron compared to the AID* degron. For PAR-6, leaky degradation was increased from 31% to 42% in the seam cells, and for SAX-7 leaky degradation increased from 45% to 72% in the ALM neuron cell body (Fig. 2B, S1B). Despite these reduced protein levels, no overt phenotypes were observed for any of these proteins. Together, these data corroborate that for some target proteins leaky degradation is a potential caveat of the AID system. In addition, the use of a more effective degron, such as mIAA7, can increase leaky degradation.

### The mIAA7 degron is compatible with the *C*.*e*.AID2 system

Recently, the AID2 system utilizing a TIR1 variant altered to fit the bulky auxin-derivative 5-Ph-IAA was adapted for *C. elegans* and shown to sharply reduce or eliminate leaky degradation while requiring low micromolar concentrations of 5-Ph-IAA (Hills-Muckey *et al*., 2022; Negishi *et al*., 2022). Given these advantages, we set out to test whether the *C*.*e*.AID2 system is compatible with the novel mIAA7 degron. We used CRISPR/Cas9 genome editing to introduce the F79G mutation into the TIR1 transgene in our PAR-6 and SAX-7 mIAA7 strains. Protein fluorescence levels were measured in these strains comparing animals not expressing TIR1, animals expressing TIR1[F79G] in the absence of auxin analog, and animals expressing TIR1[F79G] in the presence of 1 μm 5-Ph-IAA. Protein levels of PAR-6 in the seam cells and SAX-7 ALM neuron cell body were not affected by the presence of TIR1[F79G], demonstrating that the mIAA7 degron does not induce leaky degradation when combined with TIR1[F79G]. Importantly, exposure to 1 μm 5-Ph-IAA yielded complete degradation of the target proteins (Fig. 3A, B). Thus, similar to the AID* and mAID degrons that were previously tested (Hills-Muckey *et al*., 2022; Negishi *et al*., 2022), the mIAA7 degron is compatible with the AID2 TIR1[F79G] variant, resulting in effective degradation at low auxin analog concentrations and no observable leaky degradation.

**Figure 3.**
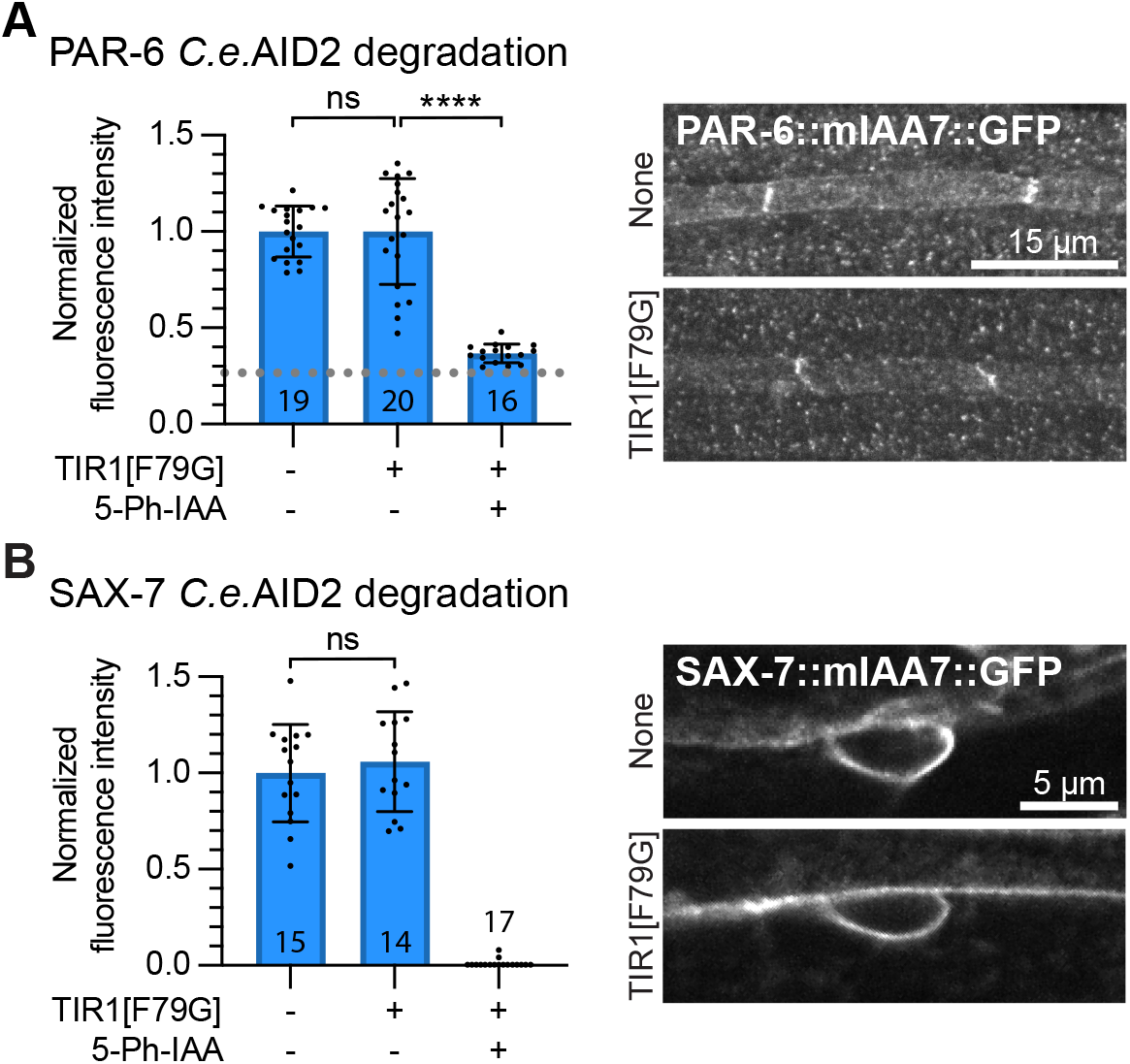
The mIAA7 degron is compatible with the *C*.*e*.AID2 system. **(A, B)** mIAA7-mediated degradation of PAR-6 in the seam cells (A) and of SAX-7 in the ALM neuron (B) of L2 larvae using the *C*.*e*.AID2 system. Values are normalized to the mean intensity levels of animals expressing the same degron tagged proteins but not expressing TIR1 and not treated with 5-Ph-IAA (*i*.*e*., the first bar). Each data point in the graph represents a single animal. Error bars: mean ± SD; Statistical test: one-way ANOVA with Šidák multiple testing correction; n values are indicated in or above the bars. Images shown are representative maximum intensity projections that were acquired and displayed with the same settings for comparison.

### The use of multiple degrons can improve the efficacy of the AID system

As a final test of the efficacy of the mIAA7 degron, we tagged the ezrin/radixin/moesin ortholog ERM-1 with mIAA7. ERM-1 is highly expressed in the intestine, where it is required for microvilli formation and patterning of the lumen. The ERM-1 protein appears highly stable and shows little turnover at the apical domain (Ramalho *et al*., 2020; Sepers *et al*., 2022). Previously, we have tested both AID and ZIF-1-mediated depletion approaches, neither of which resulted in full depletion of intestinal ERM-1 (Ramalho *et al*., 2020; Sepers *et al*., 2022). Therefore, we wanted to test if the novel mIAA7 degron could improve ERM-1 degradation. We tagged the endogenous *erm-1* locus with mCherry and mIAA7 or AID* degron coding sequences using CRISPR/Cas9 genome engineering. We then compared ERM-1 degradation in the intestine using TIR1[F79G] and 1 μm 5-Ph-IAA, measuring fluorescence levels at the apical domain of the intestine throughout larval development in synchronized animals. Whereas the AID* degron did not yield any detectable degradation, the use of the mIAA7 degron caused significant depletion of ERM-1, again confirming that mIAA7 improves the efficacy of the AID system (Fig. 4A). However, ∼76% of ERM-1 remained after 24 hours and fluorescence levels did not further decrease in the following days. Thus, mIAA7-mediated degradation of ERM-1 in the intestine is still inefficient.

**Figure 4.**
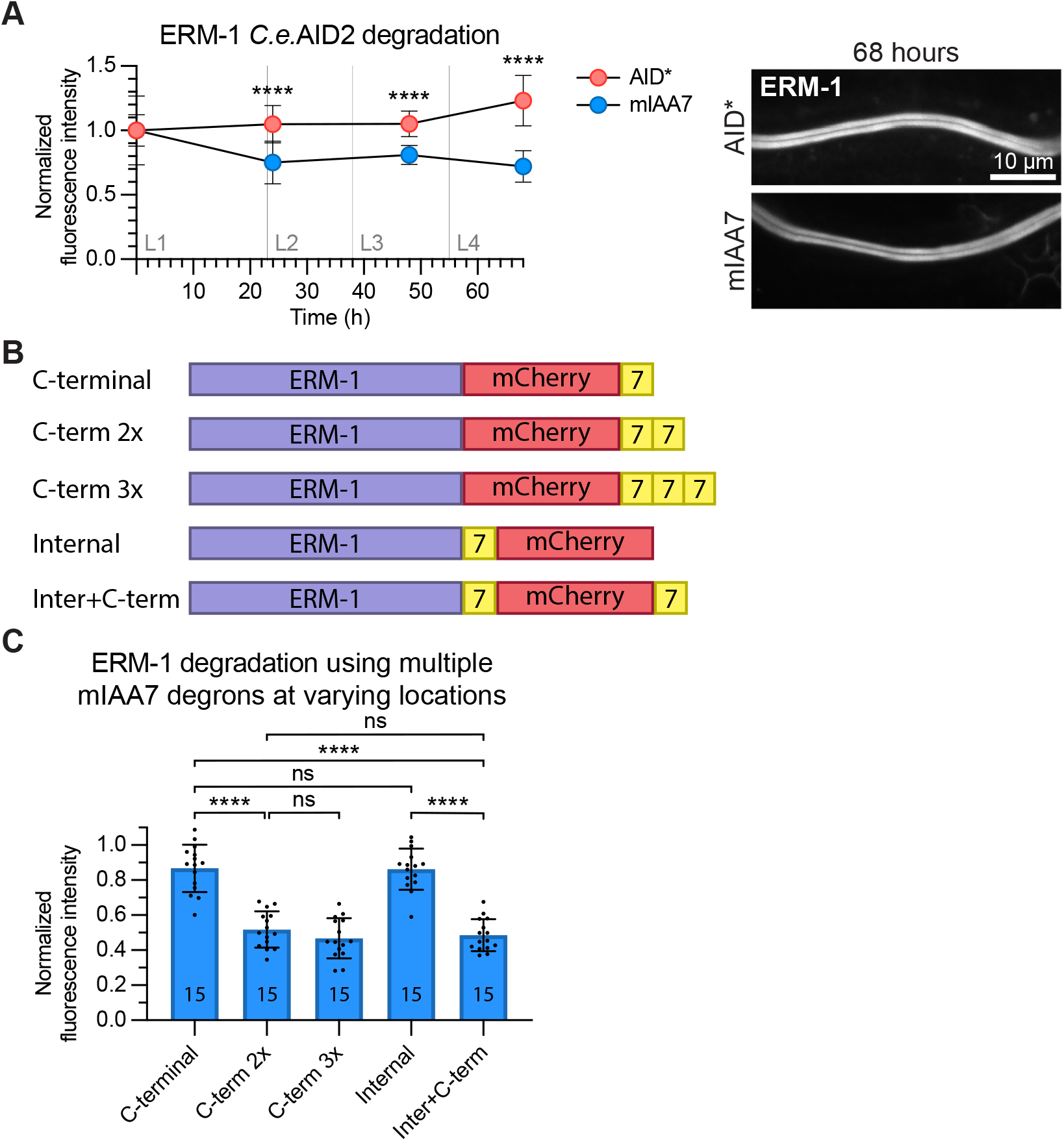
Tagging ERM-1 with multiple mIAA7 degrons improves auxin mediated degradation. **(A)** Comparison between AID*-and mIAA7-mediated degradation of intestinal ERM-1 throughout larval development using the *C*.*e*.AID2 system. Data points represent the mean intensity normalized to the mean intensity levels of animals with the same genotype and age that were not exposed to 5-Ph-IAA. Error bars: mean ± SD; Statistical test: Mann-Whitney U test; n = 12-16 animals. Images shown are representative maximum intensity projections that were acquired and displayed with the same settings for comparison. **(B)** Schematic overview of the different ERM-1 alleles used for testing the effect of multiple degrons and degron location on auxin-mediated degradation efficiency. **(C)** Comparison of intestinal degradation of ERM-1 tagged with one or multiple mIAA7 degrons at varying locations in L3 larvae using the *C*.*e*.AID2 system. Values are normalized to the mean intensity levels of animals with the same genotype that were not exposed to 5-Ph-IAA, and each data point represents a single animal. Error bars: mean ± SD; Statistical test: one-way ANOVA with Šidák multiple testing correction; n values are indicated in the bars.

In yeast, it has been shown that addition of multiple degrons to a protein can improve the AID-mediated degradation rate (Kubota *et al*., 2013; Zhang *et al*., 2022). In addition, in human cell lines the degradation rate has been shown to be affected by the localization of the degron tag (Li *et al*., 2019). We therefore investigated whether altering the number or position of degrons could improve ERM-1 degradation. Originally, ERM-1 was C-terminally tagged with mCherry followed by the mIAA7 degron (Fig. 4B - “C-terminal”). To test the effect of multiple degrons, we engineered similar alleles with double and triple mIAA7 degrons (Fig. 4B - “C-term 2x”, “C-term 3x”). To test the influence of the degron position we also generated an allele with mIAA7 located internally between the ERM-1 and mCherry coding sequences (Fig. 4B - “Internal”) and an allele combining the internal and C-terminal mIAA7 tags (Fig. 4B - “Inter+C-term”).

Synchronized animals carrying the different mIAA7 *erm-1* alleles and expressing TIR1[F79G] in the intestine were exposed to 5-Ph-IAA for 24 hours and apical ERM-1 fluorescence levels were measured. Adding an additional degron to the ERM-1 C-terminus significantly improved the degradation, depleting ∼3.5 times more protein than a single degron. Nevertheless, ∼50% of ERM-1 protein remained present. Adding a third degron to the C-terminus did not result in a further increase in degradation. We did not observe a significant difference in depletion levels between ERM-1 variants with different locations of the mIAA7 tags. Both single mIAA7 tagged alleles showed similar levels of ERM-1 depletion, as did the double ERM-1 tags “C-term 2x” and “Inter+C-term”. This suggests that the positioning of the degron does not greatly alter AID-mediated degradation efficacy. In addition, when using multiple degrons in ERM-1, it is irrelevant whether they are placed in tandem or at distinct positions. In summary, AID-mediated protein degradation of ERM-1 can be further improved by using multiple degrons tags, while the position of the degrons has no significant effect on degradation efficiency.

### Plasmid toolkit

To facilitate CRISPR/Cas9–mediated tagging of target proteins with the mIAA7 degron and commonly used fluorescent proteins we generated a plasmid toolkit (Fig. S3B). The plasmids encode one of five different fluorescent proteins flanked by the mIAA7 degron on either the N- or C-terminal side, and by a 12-amino acid glycine-rich linker on the opposing side. Using these plasmids to generate dsDNA repair templates allows for tagging of a protein of interest both N- and C-terminally with a choice of positioning the mIAA7 degron between the fluorescent protein and the protein of interest, or at the exposed terminus of the fluorescent protein. By designing primers with homology arms as overhangs according to established CRISPR-Cas9 protocols (Dokshin *et al*., 2018; Paix *et al*., 2015) this plasmid toolkit greatly convenes the generation of double-stranded DNA repair templates for CRISPR/Cas9 genome engineering of fluorescently tagged AID alleles.

## Discussion

The AID system is becoming increasingly popular in the *C. elegans* field due to its ability to degrade target proteins with both spatial and temporal control (Ashley *et al*., 2021; Hills-Muckey *et al*., 2022; Negishi *et al*., 2022; Zhang *et al*., 2015). However, the degradation efficiency varies between proteins, and not all proteins are depleted to the extent that expected phenotypes are observed (Duong *et al*., 2020; Patel and Hobert, 2017; Riga *et al*., 2021; Serrano-Saiz *et al*., 2018). Here, we demonstrated that an alternative degron sequence, termed mIAA7, improves the efficiency of degradation in *C. elegans*, as previously reported in human cell culture (Li *et al*., 2019).

The improvements to degradation efficiency appear to be applicable to most proteins, as we observed increased depletion efficiency for six proteins tested, each with different sub-cellular localizations and investigated in different tissue types. The only exception was the nuclear protein HIS-72, which was depleted with similar efficiency. In human cells, mIAA7 did not improve the depletion of the highly expressed nuclear proteins LMNA and LMNB1 (Li *et al*., 2019). However, depletion of these proteins was strongly increased when using an auxin receptor F-box protein targeted to the nucleus. A similar modification may therefore improve the depletion of nuclear proteins in *C. elegans* as well.

There are two main differences between mIAA7 and AID* that could contribute to the difference in degradation efficiency. First, mIAA7 is derived from IAA7 instead of IAA17, and hence differs in exact primary sequence. Second, the two degrons contain different parts of the IAA protein. While both degrons contain IAA Domain II, mIAA7 has a longer N-terminal extension and shorter C-terminal extension than AID* (Fig. 1A). A degron derived of IAA17 using the same protein region as mIAA7 resulted in a degradation speed close to mIAA7 in human cell culture (Li *et al*., 2019), which suggests that it is the region of the IAA protein used rather than the primary sequence that determines degradation efficiency. In support of this hypothesis, the region N-terminal to Domain II was shown to be required for TIR1 mediated degradation in plants (Dreher *et al*., 2006; Moss *et al*., 2015). However, replacing the mIAA7 sequence downstream of Domain II with the same region from IAA12 lowered the degradation speed, indicating that primary sequence can affect degradation efficiency as well (Niemeyer *et al*., 2020). One way the primary sequence could affect degradation efficiency is by providing lysine residues that can be ubiquitinated. However, the conserved KR sequence, hypothesized to be important for degradation (Dreher *et al*., 2006; Moss *et al*., 2015), is absent from the mIAA7 degron and including it did not improve degradation in human cell culture (Li *et al*., 2019). Moreover, all IAA7 ubiquitination sites that were found using mass spectrometry or are bioinformatically predicted are absent from the mIAA7 degron sequence (Niemeyer *et al*., 2020; Wang *et al*., 2021). Finally, IAA1 has been reported to be ubiquitinated and degraded without any lysine residues (Gilkerson *et al*., 2015). These data suggest that the degron itself is not ubiquitinated by the TIR1-SCF complex, or that IAA proteins can be ubiquitinated in a non-canonical manner. An alternative explanation for the effects of primary sequence on degradation efficiency is offered by a recent model based on crosslinking proteomics, in which the regions flanking Domain II contribute to the stability of the interaction between TIR1/Auxin Signaling F-Box (AFB) proteins, IAA proteins, and auxin (Niemeyer *et al*., 2020). While IAA7 showed the strongest interaction with TIR1 out of 8 IAA proteins by yeast two-hybrid (Calderón Villalobos *et al*., 2012; Niemeyer *et al*., 2020), *Arabidopsis* alone encodes 29 IAA proteins and six AFBs (Liscum and Reed, 2002; Luo *et al*., 2018; Morffy and Strader, 2022; Parry *et al*., 2009). Therefore, even more effective IAA-AFB combinations may be discovered.

One of the drawbacks of the AID system is the potential for degradation in the absence of auxin. In our experiments, RPS-26 and BBLN-1 showed little leaky degradation with either degron, but leaky degradation for PAR-6 and SAX-7 was relatively high for AID* and was enhanced by use of the mIAA7 degron. Importantly, when combining the mIAA7 degron with the *C*.*e*.AID2 system that utilizes TIR1[F79G] and the auxin-derivative 5-Ph-IAA, we observed no leaky degradation of PAR-6 and SAX-7 in the absence of auxin/5-Ph-IAA. Thus, the benefits of the mIAA7 degron can be realized without the drawback of increased leaky degradation. Additional measures that users could take to address leaky degradation include reducing the expression levels of TIR1, using AFB2 instead of TIR1 (Li *et al*., 2019), or re-engineering the degron sequence to include the PB1 domain of Auxin Response Factor plant proteins, which was shown to reduce leaky degradation of AID degron tagged proteins in human cell culture (Sathyan *et al*., 2019).

Despite the improvements in degradation kinetics conferred by the mIAA7 degron, some proteins remain refractive to full depletion. For difficult to degrade proteins, the addition of multiple degron sequence can further increase the degradation efficiency. In our hands, auxin-mediated degradation of ERM-1 was improved when tagged with two mIAA7 degrons compared to a single degron. This finding is in line with previous results in yeast, in which using multiple degrons improved degradation (Kubota *et al*., 2013; Zhang *et al*., 2022). However, in contrast to results in yeast, we did not observe a further improvement in the depletion efficiency when using three degrons. This difference may be specific to ERM-1, which associates very stably with the apical domain of intestinal cells (Ramalho *et al*., 2020; Sepers *et al*., 2022). It is possible, therefore, that a fraction of ERM-1 is not accessible to the SCF complex or the proteasome and that all accessible ERM-1 is degraded after 24 hours using two degrons. Alternatively, the ubiquitination or degradation machinery may be rate limiting due to the high expression levels of ERM-1.

Interestingly, the position of the degrons did not seem to affect the degradation rate of ERM-1. Two degrons in tandem resulted in similar depletion levels as two degrons flanking the mCherry sequence on both sides. We did not test an N-terminal placement of the degron as the two predicted splice variants of ERM-1 do not share the same N-terminus, resulting in the degradation of only a subpopulation of ERM-1. Our results contrast with data from human cell culture in which degron position did influence the degradation rate for two proteins tested (Li *et al*., 2019). For the protein SEC61B degradation was higher when mIAA7 was positioned at the extreme N-terminus compared to an internal position between a fluorescent protein and SEC61B. For the protein Seipin, a C-terminal degron performed better than an N-terminal degron, and including a fluorescent protein downstream of the C-terminal degron further increased degradation efficiency. Given the low number of proteins tested in *C. elegans* and human cells, the exact influence of degron position requires further investigation.

In summary, the mIAA7 degron further extends the usability of the AID system in *C. elegans*, improving degradation efficiency while, particularly in combination with *C*.*e*.AID2, not affecting the steady-state level of proteins in the absence of auxin.

## Methods

### *C. elegans* Strains and Culture Conditions

*C. elegans* strains were cultured under standard conditions (Brenner, 1974). Only hermaphrodites were used. All experiments were performed with animals grown at 15 °C or 20 °C on standard Nematode Growth Medium (NGM) agar plates seeded with OP50 *Escherichia coli*. The HIS-72 timecourse experiment was performed in liquid culture (see below). Table 1 contains a list of all the strains used and their source.

**Table 1.**
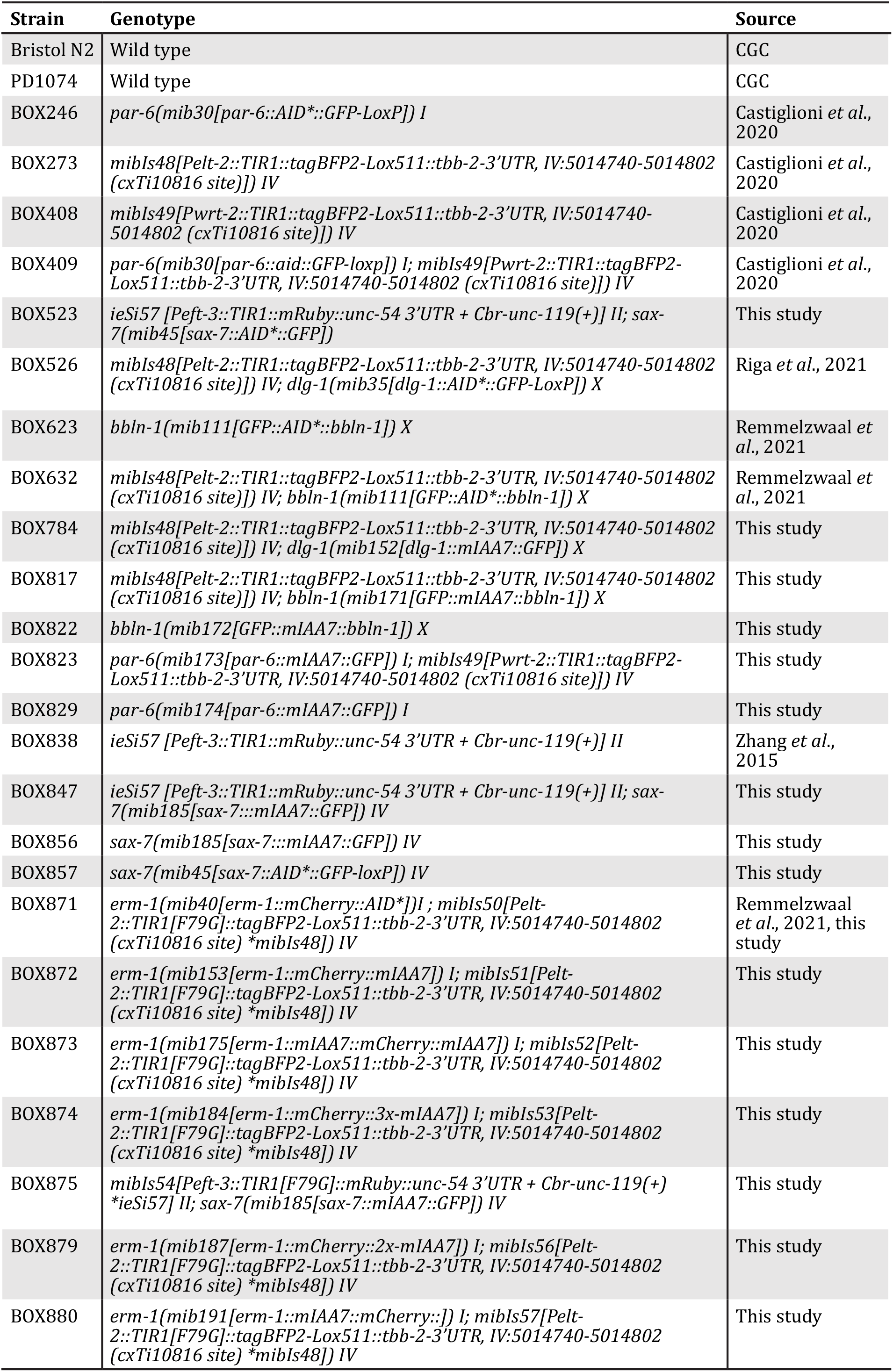
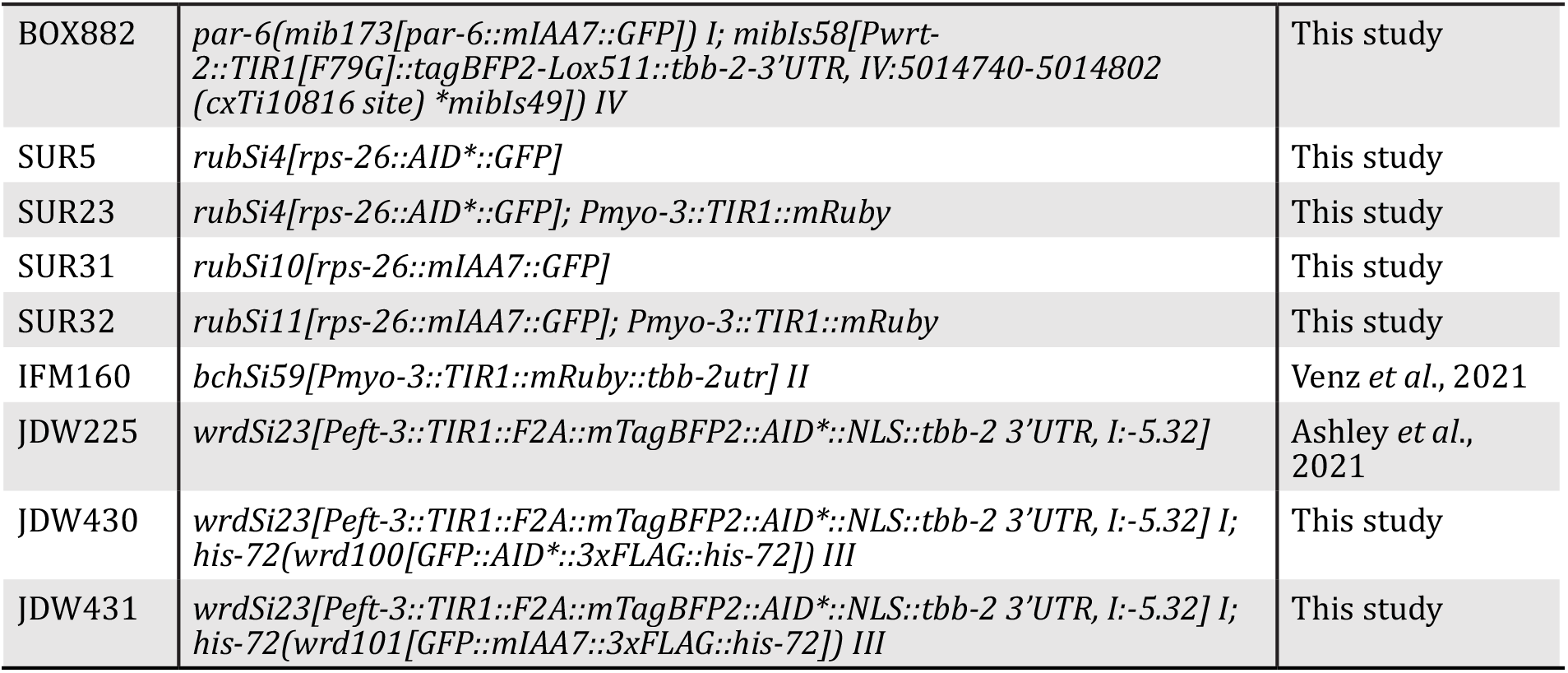
List of strains used.

| Strain | Genotype | Source |
| --- | --- | --- |
| Bristol N2 | Wild type | CGC |
| PD1074 | Wild type | CGC |
| BOX246 | *par-6(mib30[par-6::AID\*::GFP-LoxP]) I* | Castiglioni *et al.*, 2020 |
| BOX273 | *mibIs48[Pelt-2::TIR1::tagBFP2-Lox511::tbb-2-3'UTR, IV:5014740-5014802 (cxTi10816 site)] IV* | Castiglioni *et al.*, 2020 |
| BOX408 | *mibIs49[Pwrt-2::TIR1::tagBFP2-Lox511::tbb-2-3'UTR, IV:5014740-5014802 (cxTi10816 site)] IV* | Castiglioni *et al.*, 2020 |
| BOX409 | *par-6(mib30[par-6::aid::GFP-loxp]) I; mibIs49[Pwrt-2::TIR1::tagBFP2-Lox511::tbb-2-3'UTR, IV:5014740-5014802 (cxTi10816 site)] IV* | Castiglioni *et al.*, 2020 |
| BOX523 | *ieSi57 [Peft-3::TIR1::mRuby::unc-54 3'UTR + Cbr-unc-119(+)] II; sax-7(mib45[sax-7::AID\*::GFP])* | This study |
| BOX526 | *mibIs48[Pelt-2::TIR1::tagBFP2-Lox511::tbb-2-3'UTR, IV:5014740-5014802 (cxTi10816 site)] IV; dlg-1(mib35[dlg-1::AID\*::GFP-LoxP]) X* | Riga *et al.*, 2021 |
| BOX623 | *bbln-1(mib111[GFP::AID\*::bbln-1]) X* | Rommelzwaal *et al.*, 2021 |
| BOX632 | *mibIs48[Pelt-2::TIR1::tagBFP2-Lox511::tbb-2-3'UTR, IV:5014740-5014802 (cxTi10816 site)] IV; bbln-1(mib111[GFP::AID\*::bbln-1]) X* | Rommelzwaal *et al.*, 2021 |
| BOX784 | *mibIs48[Pelt-2::TIR1::tagBFP2-Lox511::tbb-2-3'UTR, IV:5014740-5014802 (cxTi10816 site)] IV; dlg-1(mib152[dlg-1::mIAA7::GFP]) X* | This study |
| BOX817 | *mibIs48[Pelt-2::TIR1::tagBFP2-Lox511::tbb-2-3'UTR, IV:5014740-5014802 (cxTi10816 site)] IV; bbln-1(mib171[GFP::mIAA7::bbln-1]) X* | This study |
| BOX822 | *bbln-1(mib172[GFP::mIAA7::bbln-1]) X* | This study |
| BOX823 | *par-6(mib173[par-6::mIAA7::GFP]) I; mibIs49[Pwrt-2::TIR1::tagBFP2-Lox511::tbb-2-3'UTR, IV:5014740-5014802 (cxTi10816 site)] IV* | This study |
| BOX829 | *par-6(mib174[par-6::mIAA7::GFP]) I* | This study |
| BOX838 | *ieSi57 [Peft-3::TIR1::mRuby::unc-54 3'UTR + Cbr-unc-119(+)] II* | Zhang *et al.*, 2015 |
| BOX847 | *ieSi57 [Peft-3::TIR1::mRuby::unc-54 3'UTR + Cbr-unc-119(+)] II; sax-7(mib185[sax-7::mIAA7::GFP]) IV* | This study |
| BOX856 | *sax-7(mib185[sax-7::mIAA7::GFP]) IV* | This study |
| BOX857 | *sax-7(mib45[sax-7::AID\*::GFP-loxP]) IV* | This study |
| BOX871 | *erm-1(mib40[erm-1::mCherry::AID\*]) I; mibIs50[Pelt-2::TIR1[F79G]::tagBFP2-Lox511::tbb-2-3'UTR, IV:5014740-5014802 (cxTi10816 site) \*mibIs48]) IV* | Rommelzwaal *et al.*, 2021, this study |
| BOX872 | *erm-1(mib153[erm-1::mCherry::mIAA7]) I; mibIs51[Pelt-2::TIR1[F79G]::tagBFP2-Lox511::tbb-2-3'UTR, IV:5014740-5014802 (cxTi10816 site) \*mibIs48]) IV* | This study |
| BOX873 | *erm-1(mib175[erm-1::mIAA7::mCherry::mIAA7]) I; mibIs52[Pelt-2::TIR1[F79G]::tagBFP2-Lox511::tbb-2-3'UTR, IV:5014740-5014802 (cxTi10816 site) \*mibIs48]) IV* | This study |
| BOX874 | *erm-1(mib184[erm-1::mCherry::3x-mIAA7]) I; mibIs53[Pelt-2::TIR1[F79G]::tagBFP2-Lox511::tbb-2-3'UTR, IV:5014740-5014802 (cxTi10816 site) \*mibIs48]) IV* | This study |
| BOX875 | *mibIs54[Peft-3::TIR1[F79G]::mRuby::unc-54 3'UTR + Cbr-unc-119(+)] \*ieSi57] II; sax-7(mib185[sax-7::mIAA7::GFP]) IV* | This study |
| BOX879 | *erm-1(mib187[erm-1::mCherry::2x-mIAA7]) I; mibIs56[Pelt-2::TIR1[F79G]::tagBFP2-Lox511::tbb-2-3'UTR, IV:5014740-5014802 (cxTi10816 site) \*mibIs48]) IV* | This study |
| BOX880 | *erm-1(mib191[erm-1::mIAA7::mCherry::]) I; mibIs57[Pelt-2::TIR1[F79G]::tagBFP2-Lox511::tbb-2-3'UTR, IV:5014740-5014802 (cxTi10816 site) \*mibIs48]) IV* | This study |
| BOX882 | *par-6(mib173[par-6::mIAA7::GFP]) I; mibls58[Pwrt-2::TIR1[F79G]::tagBFP2-Lox511::tbb-2-3'UTR, IV:5014740-5014802 (cxTi10816 site) \*mibls49]) IV* | This study |
| SUR5 | *rubSi4[rps-26::AID\*::GFP]* | This study |
| SUR23 | *rubSi4[rps-26::AID\*::GFP]; Pmyo-3::TIR1::mRuby* | This study |
| SUR31 | *rubSi10[rps-26::mIAA7::GFP]* | This study |
| SUR32 | *rubSi11[rps-26::mIAA7::GFP]; Pmyo-3::TIR1::mRuby* | This study |
| IFM160 | *bchSi59[Pmyo-3::TIR1::mRuby::tbb-2utr] II* | Venz *et al.*, 2021 |
| JDW225 | *wrdSi23[Peft-3::TIR1::F2A::mTagBFP2::AID\*::NLS::tbb-2 3'UTR, I:-5.32]* | Ashley *et al.*, 2021 |
| JDW430 | *wrdSi23[Peft-3::TIR1::F2A::mTagBFP2::AID\*::NLS::tbb-2 3'UTR, I:-5.32] I; his-72(wrd100[GFP::AID\*::3xFLAG::his-72]) III* | This study |
| JDW431 | *wrdSi23[Peft-3::TIR1::F2A::mTagBFP2::AID\*::NLS::tbb-2 3'UTR, I:-5.32] I; his-72(wrd101[GFP::mIAA7::3xFLAG::his-72]) III* | This study |

### Cloning of the mIAA7 repair template plasmids

The mIAA7 repair templates were cloned into pBSK+ (Addgene #212207) using the Gibson assembly cloning strategy (Gibson *et al*., 2009). Every DNA fragment was amplified, and plasmid was PCR-linearized by Q5 polymerase (NEB) using primers (IDT) containing overhangs with the appropriate Gibson assembly sequences and, if applicable, the 12-aminoacid glycine-rich linker. The mIAA7 sequence, based on Li *et al*., 2019, was codon-optimized for *C. elegans* with a synthetic intron and ordered as a gBlocks Gene Fragment (IDT). The pJJS001–pJJS004 plasmids were assembled by inserting the mIAA7 and GFP or mCherry sequences directly into pBSK+. The pJJS005–pJJS012 plasmids were assembled by substituting the GFP of pJJS001 and pJJS002 plasmids with the respective fluorescent protein sequences. All assembly products were verified by Sanger sequencing (Macrogen Europe). pJW2341 was assembled by Gibson cloning a mIAA7 fragment into pJW2086 (Ashley *et al*., 2021). Cloning details are available upon request. Table 2 contains a list of all plasmids made with the respective primer sequences and origin of the fluorescent protein sequence, and full DNA sequences of all plasmids are provided in Supplemental File 1. All plasmids will be deposited to Addgene.

**Table 2.**
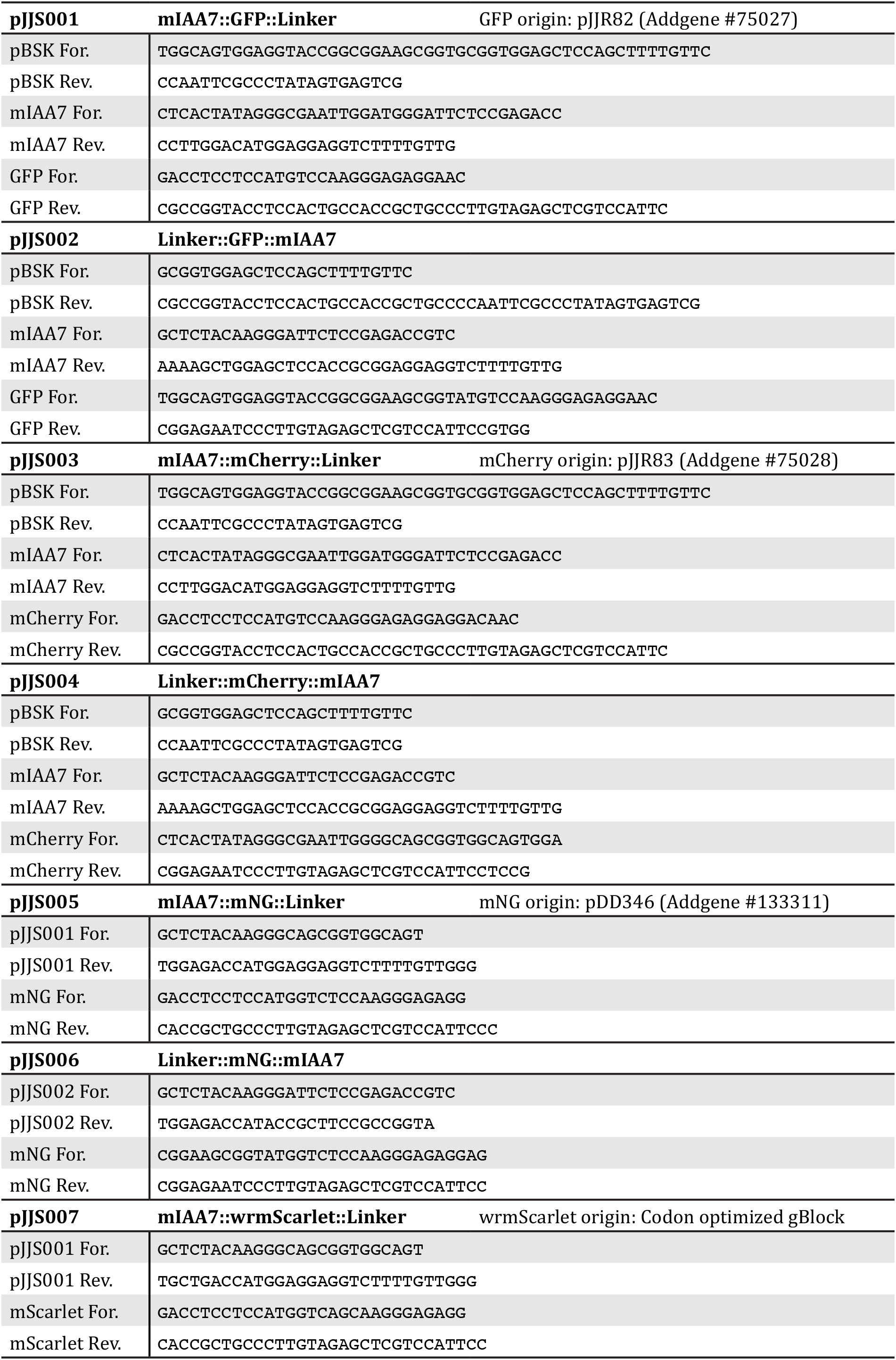
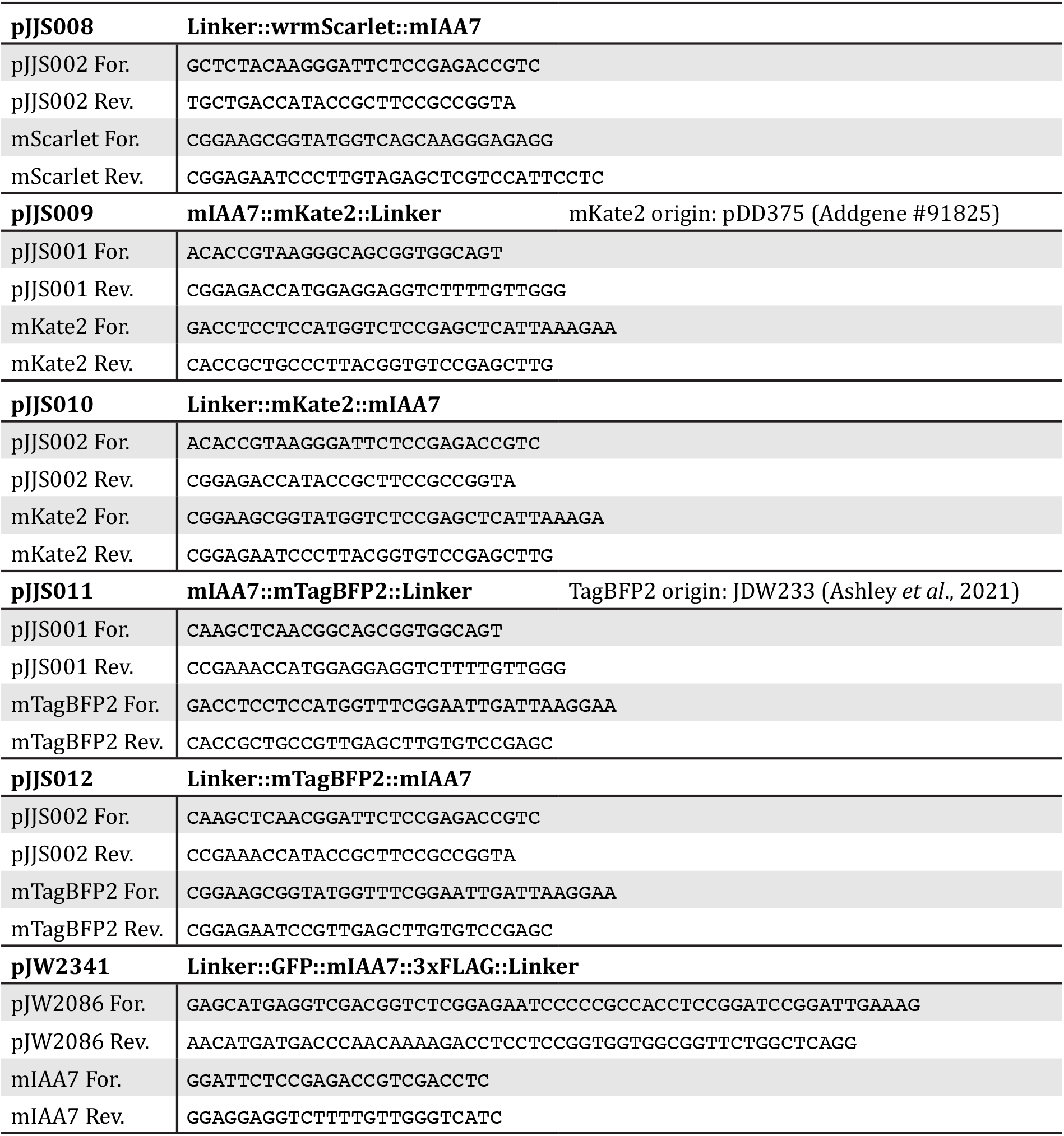
Primer sequences for plasmids.

### CRISPR/Cas9 genome editing

All alleles were made using the homology-directed repair of the CRISPR-Cas9-induced DNA double strand breaks. Repair templates included, when appropriate, silent mutations to prevent recutting of repaired loci by Cas9. To verify the edits, the insertion sites were PCR amplified and sequenced by Sanger sequencing. Table 3 contains a list of all CRISPR-Cas9 edits made with the respective primer sequences, and Supplementary file 1 contains all final genomic sequences.

**Table 3.**
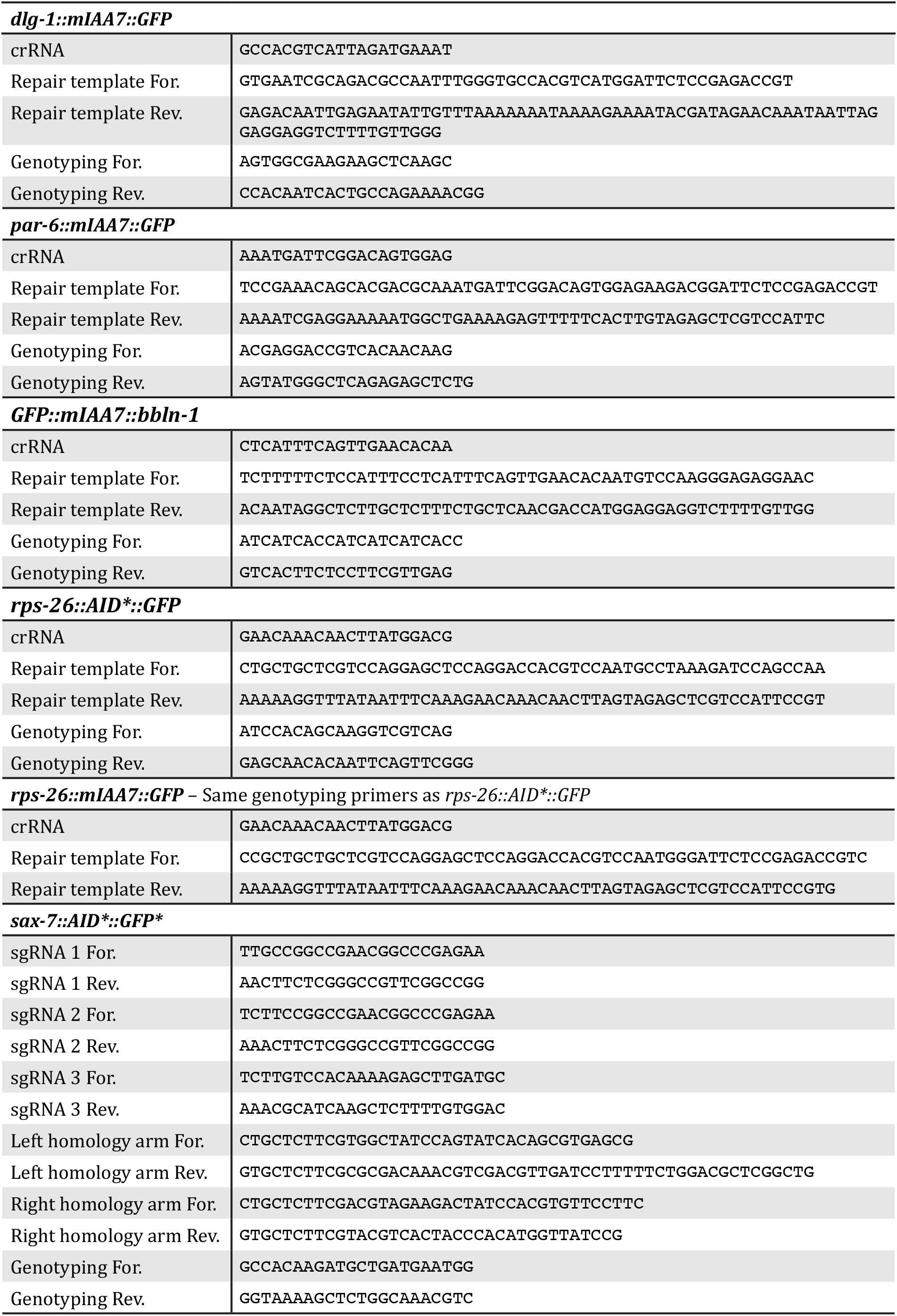
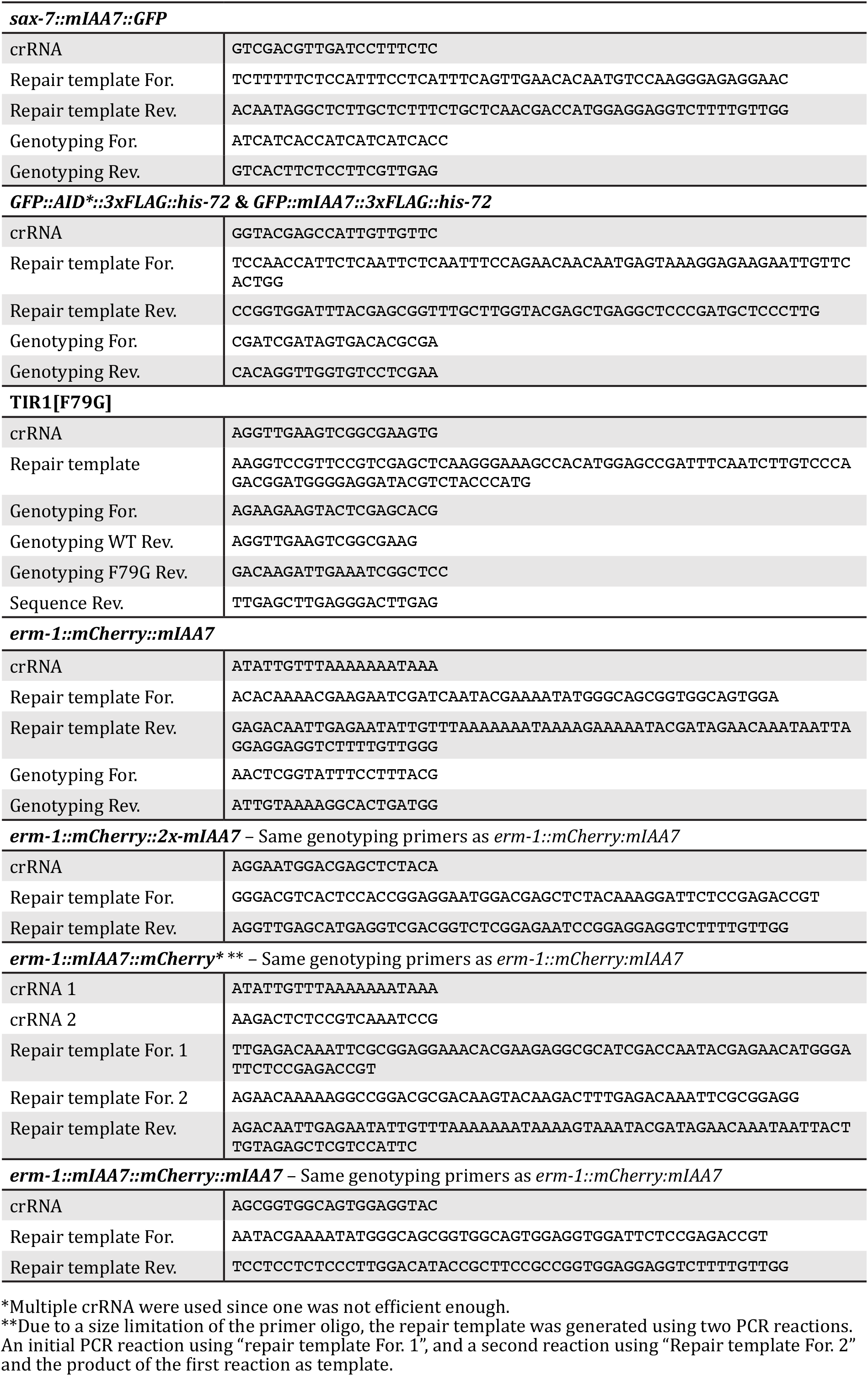
Genome engineering reagents.

The sax-7::AID*::GFP allele was made using plasmid based expression of Cas9 and sgRNA and a plasmid repair template contain a self-excising cassette for selection of candidate integrants (Dickinson *et al*., 2015). Repair template and sgRNA plasmid were designed and assembled using the SapTrap cloning strategy (Dickinson *et al*., 2015; Schwartz and Jorgensen, 2016). Reagents were injected and knock-in animals were recovered as previously descripted.

All other alleles were generated using microinjected Cas9 ribonucleoprotein complexes and linear repair templates, similar to the approach described by (Ghanta *et al*., 2021). We used a Cas9 amount of 250–700 ng/μl, a Cas9/crRNA ratio of 3.0–4.5, and a pSEM229 (Pmlc-1::mNeonGreen) or pRF4 (*rol-6(su1006)*) co-injection marker (El Mouridi *et al*., 2020; Mello *et al*., 1991). For tagging endogenous loci with a single AID* or mIAA7 degron and fluorescent protein, dsDNA repair templates were amplified using primers with 5’ SP9 modifications (IDT) from the pJRK86 plasmid (AID*::GFP, Addgene #173743) and mIAA7 repair template plasmids in table 1. For *his-72* editing, GFP::AID*::3xFLAG repair templates were generated by PCR using a pJW2086 template and GFP::mIAA7::3xFLAG repair templates were generated by PCR using a pJW2341 repair template (Table 1). For tagging ERM-1 with two degrons, a second mIAA7 degron was inserted into the existing ERM-1::mCherry::mIAA7 allele using similar dsDNA repair templates amplified from an mIAA7 repair template plasmids in table 1. The C-term 3x ERM-1 allele was generated using the same reagents as the C-term 2x allele but was the result of a fortuitous incorrect repair event. The TIR1[F79G] alleles were generated using a ssDNA oligo repair template (IDT).

### Auxin treatment of synchronized worms

Animals were developmentally synchronized by hypochlorite bleaching of gravid adults to release embryos, which were hatched overnight in M9 buffer. For HIS-72, depletion experiments were performed in liquid culture. Approximately 14,000 synchronized L1 animals were resuspended in 6 ml of M9 + 0.025% gelatin supplemented with 13.3% (v/v) concentrated OP50 and either 4% ethanol (control) or 4mM indole-3-acetic acid (IAA or auxin) in ethanol. All other depletion experiments were performed on agar plates. Synchronized L1 larvae were first placed on NGM plates seeded with *E. coli* OP50 and allowed to develop for 24–48 hours. Animals were then transferred to NGM plates seeded with *E. coli* OP50 and containing 1 mM, 50 μM, or 5 μM IAA, or 1 μM 5-Ph-IAA. Auxin NGM plates were prepared by diluting 1M IAA (Alfa Aesar) or 1mM 5-Ph-IAA (BioAcademia) dissolved in 100% ethanol into NGM agar that was cooled down to ∼50 °C prior to plate pouring.

### Western blot analysis

At each timepoint, 500 μl of animals in solution was removed (∼1160 larvae), and samples for western blotting were generated similar to (Vo *et al*., 2021), except samples were resuspended in 100 μl of M9 + 0.025% gelatin and freeze-cracked twice on dry ice before Laemmli sample buffer was added to 1X and samples were boiled for 5 minutes. Five μl of lysate was resolved on precast 4–20% Mini-Protean TGX Stain Free Gels (Bio-Rad) before being transferred to a polyvinylidene difluoride membrane by semi-dry transfer with a TransBlot Turbo (Bio-Rad). Blots were probed with 1:2000 horseradish peroxidase (HRP)-conjugated anti-FLAG M2 (Sigma-Aldrich, A8592-5×1MG, Lot #SLCB9703) or 1:2000 mouse anti-alpha-Tubulin 12G10 concentrated supernatant (Developmental Studies Hybridoma Bank). For the anti-alpha tubulin blot we used 1:10,000 Digital anti-mouse HRP conjugate (Kindle Biosciences LLC, R1005) secondary antibody. Blots were developed similar to Johnson *et al*., 2021 using Supersignal West Femto Maximum Sensitivity Substrate (Thermo Fisher Scientific, 34095) and the high-resolution setting on a ChemiDoc MP system (Bio-Rad).

### Epifluorescence microscopy

At each HIS-72 depletion timepoint images were collected by removing 100 μl of culture of the indicated genotype and treatment, adding 1 ml of M9 + 0.025% gelatin, pelleting at 700g, washing twice in M9 + 0.025% gelatin, and then transferring animals to a 2% agarose pad. 10 μl of 5 mM levamisole was added to immobilize animals and a cover slip was placed on top. Images were acquired on an AxioImager M2 microscope (Carl Zeiss Microscopy, LLC) equipped with a Colibri 7 LED light source and an Axiocam 506 mono camera using a Plan-Apochromat 63x/1.40 Oil DIC lens. Acquired images were processed through Fiji software version: 2.3.0/1.53q (Schindlin *et al*., 2012).

### Spinning disk confocal microscopy

Larvae were mounted on a 5 % agarose pad in 20 mM Tetramisole solution in M9 to induce paralysis. Spinning disk confocal imaging was performed using a Nikon Ti-U microscope equipped with a Yokogawa CSU-X1 spinning disk using a 60×-1.4 NA objective, 488 nm and 561 nm lasers, Semrock “530” GFP-L and “600” TxRed emission filters, and an Andor iXON DU-885 camera. Spinning disk images were acquired using MetaMorph Microscopy Automation & Image Analysis Software. Image scales were calibrated using a micrometer slide. All stacks along the z-axis were obtained at 0.25 μm intervals, and maximum intensity Z projections were done in Fiji software. For display in figures, level adjustments, false coloring, and image overlays were done in Adobe Photoshop. Image rotation, cropping, and panel assembly were done in Adobe Illustrator. All edits were done non-destructively using adjustment layers and clipping masks, and images were kept in their original capture bit depth until final export from Illustrator for publication.

### Quantitative image analysis

Quantitative analysis of spinning disk images was done in Fiji and Python. All measurement values were first corrected for imaging system background levels by subtracting the average of a region within the field-of-view that did not contain any animals. For the measurements of the intestinal apical junctions (DLG-1) and membrane (BBLN-1 and ERM-1), analyses were done on intestinal cells forming int2 through int6, at the center of the intestinal lumen. A maximum projection of 5 slices was made, and three 25 px-wide line scans perpendicular to the apical junction or membrane were taken per animal. For the measurements of the cortex of the cell body of the ALM neuron (SAX-7), two 10 px-wide line scans using a maximum projection of 3 slices were taken perpendicular to the membrane per animal. For each line scan, the peak value and cytoplasmic value were determined using the peak finding tools of the Scipy Signal Python datapack with rel_ height of 1.0 (Virtanen *et al*., 2020). Cytoplasmic values were subtracted from the corresponding peak values, and the resulting final junctional or membrane intensity values were averaged per animal. For SAX-7, exposure to auxin of 3 hrs or longer resulted in complete depletion and an inability to locate the ALM cell body. For these timepoints, values were plotted at 0 and no statistics were performed. For the measurement of PAR-6 at the apical domain of the seam cells and RPS-26 in the body wall muscle cytoplasm, averages of three regions were determined using a max projection of 3 slices (PAR-6) or a single slice (RPS-26).

### Statistical analysis

All statistical analyses were performed using GraphPad Prism 8. For population comparisons, a D’Agostino & Pearson test of normality was first performed to determine if the data was sampled from a Gaussian distribution. For data drawn from a Gaussian distribution, comparisons between two populations were done using an unpaired t-test, with Welch’s correction if the SDs of the populations differ significantly, and comparisons between >2 populations were done using a one-way ANOVA if the SDs of the populations differ significantly. For data not drawn from a Gaussian distribution, a non-parametric test was used (Mann-Whitney for 2 populations and Kruskal-Wallis for >2 populations). ANOVA and non-parametric tests were followed up with multiple comparison tests of significance (Dunnett’s, Tukey’s, Dunnett’s T3 or Dunn’s). Tests of significance used and sample sizes are indicated in the figure legends. No statistical method was used to pre-determine sample sizes. No samples or animals were excluded from analysis. The experiments were not randomized, and the investigators were not blinded to allocation during experiments and outcome assessment. All presented graphs were made using GraphPad Prism and Adobe Illustrator. ns is P > 0.05, * is P ≤ 0.05, ** is P ≤ 0.01, *** is P ≤ 0.001, and **** is P ≤ 0.0001.

## Acknowledgements

We thank members of the S. van den Heuvel and M. Boxem groups for helpful discussions. We thank A. Riga for providing BOX523 and BOX857, D.J. Dolfing for providing SUR5 and SUR23, and V.C. Portegijs for providing the codon optimized wrmScarlet gBlock. We also thank Wormbase (Harris et al., 2020) and the Biology Imaging Center, Faculty of Sciences, Department of Biology, Utrecht University. Some strains were provided by the Caenorhabditis Genetics Center, which is funded by NIH Office of Research Infrastructure Programs (P40 OD010440). This work was supported by the NIH/National Institute of General Medical Sciences R01 GM138701 to J.D. Ward and by the Netherlands Organization for Scientific Research (NWO) 016.VICI.170.165 grant to M. Boxem.

## Author contributions

Conceptualization: J.J.S., M.B.; Formal analysis: J.J.S., N.H.M.V., J.D.W.; Investigation: J.J.S., N.H.M.V., A.A.V., S.R., J.D.W.; Resources: J.J.S., N.H.M.V., J.M.R., S.R., J.D.W., M.B.; Writing - original draft: J.J.S., N.H.M.V.; Writing – review and editing: S.R., J.D.W., M.B.; Visualization: J.J.S.; Supervision: J.J.S., J.D.W., M.B.; Project administration: J.D.W., M.B.; Funding acquisition: J.D.W., M.B.

## Supplementary figures

**Figure S1.**
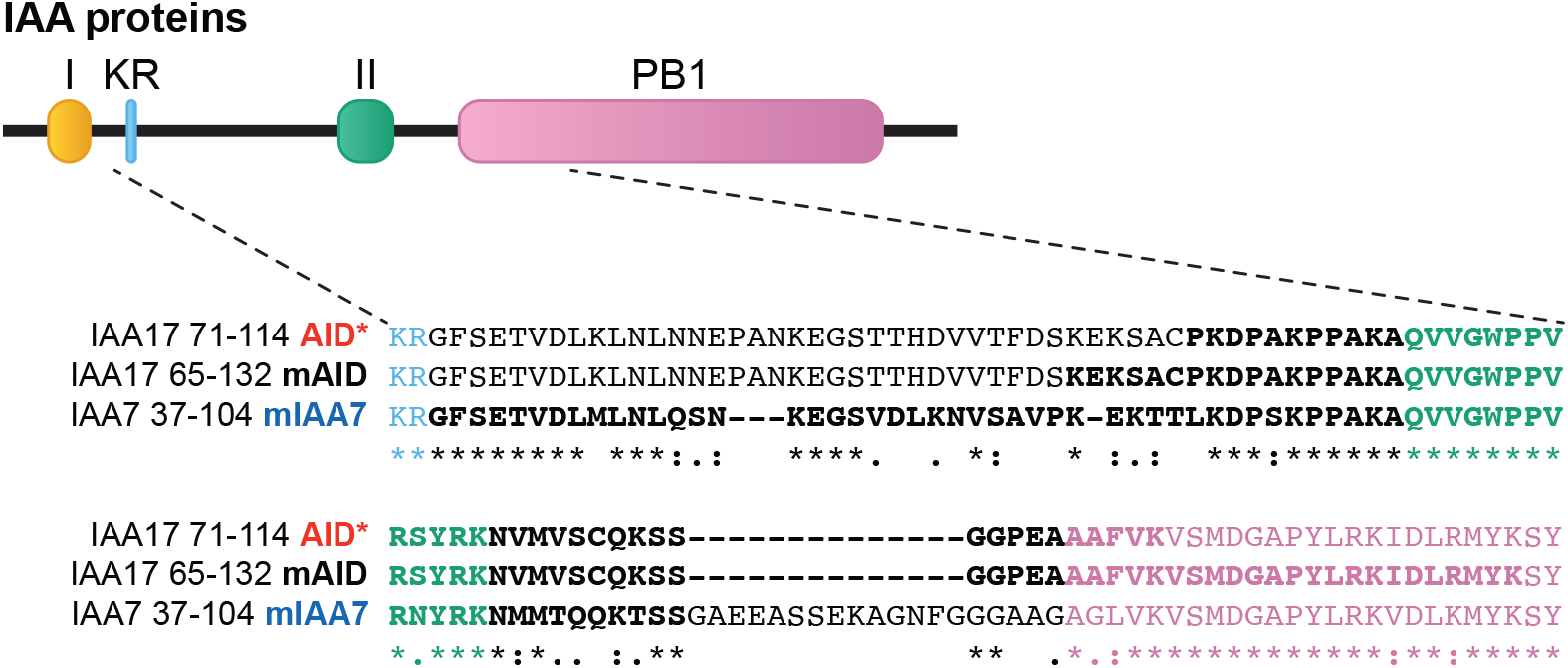
Schematic overview of the IAA proteins and the sequences of the different AID degrons that have been derived from them. Amino acids in bold are part of the degron sequences. IAA = Indole-3-acetic acid; AID = auxin inducible degron; I = domain I; KR = conserved lysine and arginine residue; II = domain II; PB1 = Phox and Bem1p domain.

**Figure S2.**
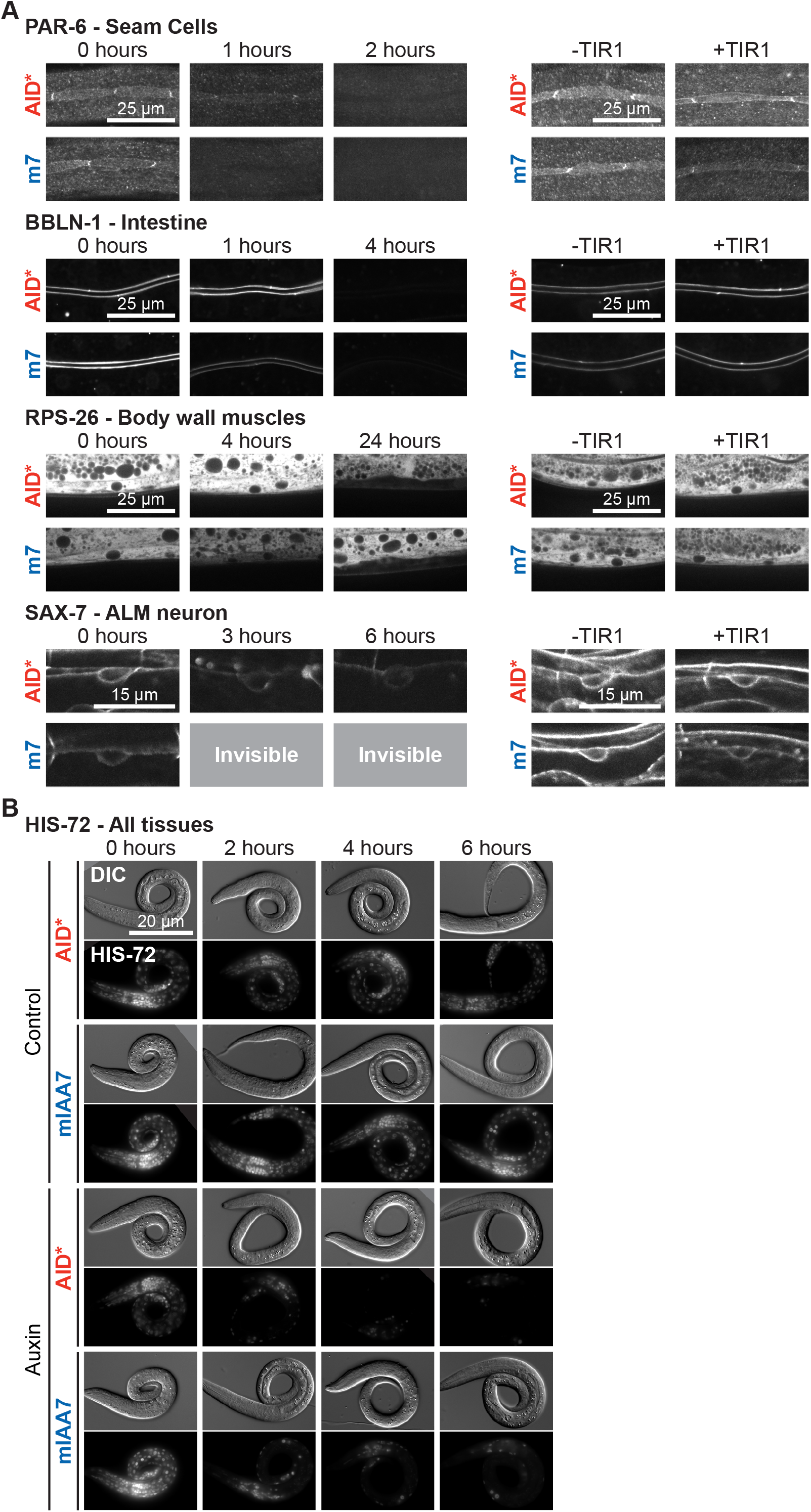
The mIAA7 degron robustly increases the efficiency of AID-induced protein degradation for several proteins across multiple tissues and cellular compartments. **(A)** Comparison between AID*- and mIAA7-mediated degradation for indicated proteins and tissues. PAR-6 was measured at the apical domain of seam cells in L2 larvae on 5 μM auxin, BBLN-1 was measured at the apical domain in the intestine of L3 larvae treated with 50 μM auxin, SAX-7 was measured at the plasma membrane in the ALM neuron cell body of L3 larvae treated with 5 μM auxin, and RPS-26 was measured in the cytoplasm of body wall muscles of L2 larvae treated with 1 mM auxin. m7 = mIAA7 **(B)** Comparison between AID*-and mIAA7-mediated degradation for HIS-72 in all tissues of synchronized control or 4 mM auxin-treated L1 larvae. Images shown are representative maximum intensity projections that were acquired and displayed with the same settings for comparison, except for RPS-26 and HIS-72 for which a single plane is presented.

**Figure S3.**
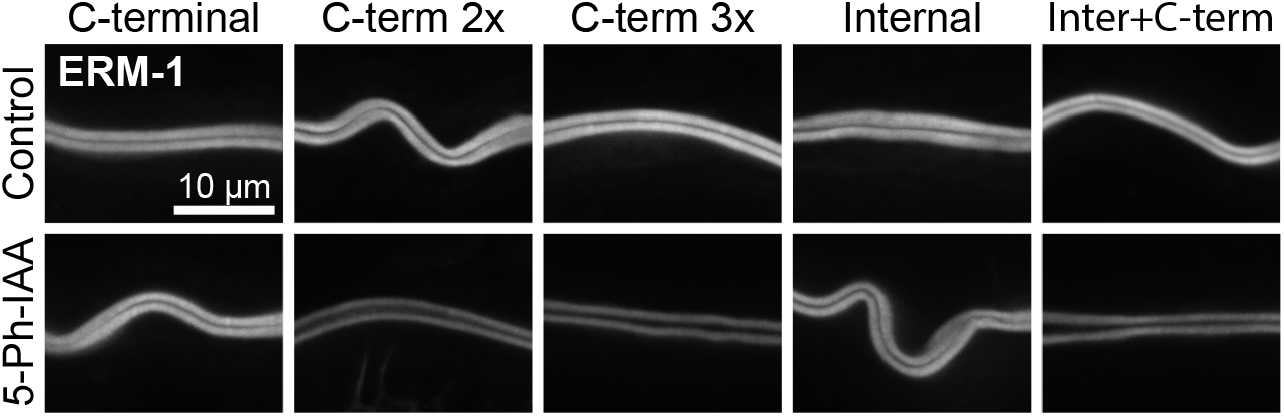
Tagging proteins with multiple mIAA7 degrons can improve auxin mediated degradation. Comparison of intestinal degradation of ERM-1 tagged at indicated locations with one or multiple mIAA7 degrons in L3 larvae using the *C*.*e*.AID2 system. Images shown are representative maximum intensity projections that were acquired and displayed with the same settings for comparison.

**Figure S4.**
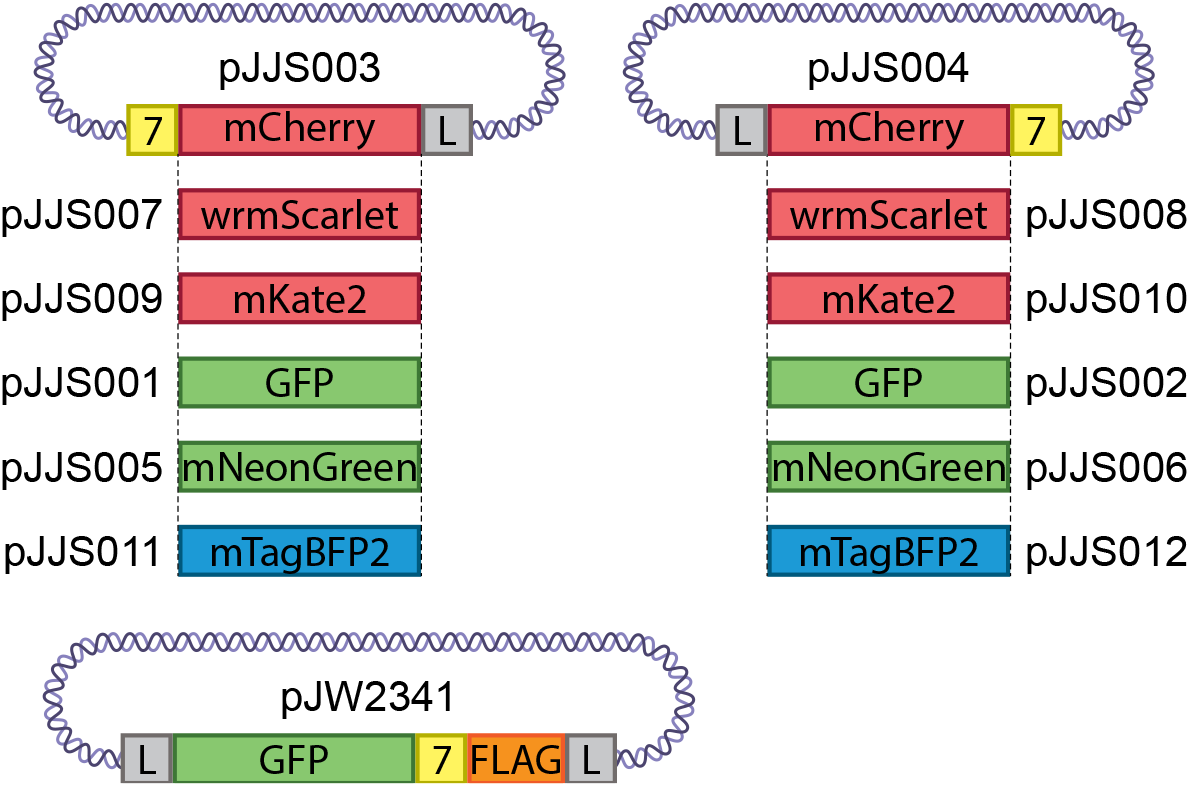
Schematic overview of the mIAA7 repair template plasmids. 7 = mIAA7 degron; L = linker sequence; GFP = green fluorescent protein; BFP = blue fluorescent protein; FLAG = 3x FLAG-tag.

